# Recovered and dead outcome patients caused by influenza A (H7N9) virus infection show different pro-inflammatory cytokine dynamics during disease progress and its application in real-time prognosis

**DOI:** 10.1101/339333

**Authors:** Yingxia Liu, Xinfa Wang, Houshun Zhu, Jinmin Ma, Zhe Lu, Jing Yuan, Jianming Li, Jiandong Li, Yan Ren, Bo Wen, Wenjie Ouyang, Haixia Zheng, Rongrong Zou, Yuhai Bi, Changcheng Yin, Zhenyu Guo, Wanying Sun, Na Pei, Junhua Li, Shida Zhu, Huanming Yang, Lei Liu, Xun Xu, Siqi Liu, Hui Wang, Liqiang Li

**Affiliations:** Shenzhen Key Laboratory of Pathogen and Immunity, State Key Discipline of Infectious Disease, Shenzhen Third People’s Hospital, Shenzhen 518112, China; BGI-Shenzhen, Shenzhen 518083, China; China National Genebank, BGI-Shenzhen, Shenzhen 518083, Guangdong, China; BGI Education Center, University of Chinese Academy of Sciences, Shenzhen 518083, Guangdong, China; Qingdao University BGI Joint Innovation College, Qingdao University, Qingdao 266071, China; CAS Key Laboratory of Pathogenic Microbiology and Immunology, Collaborative Innovation Center for Diagnosis and Treatment of Infectious Disease, Institute of Microbiology, Center for Influenza Research and Early-warning (CASCIRE), Chinese Academy of Sciences, Beijing 100101, China; Beijing Protein Innovation Co. Ltd, Beijing, China; Department of Engineering Science, University of Oxford, Oxford OX3 7DQ, UK; James D. Watson Institute of Genome Science, Hangzhou, China

## Abstract

The persistent circulation of influenza A(H7N9) virus within poultry markets and human society leads to sporadic epidemics of influenza infections. Severe pneumonia and acute respiratory distress syndrome (ARDS) caused by the virus lead to high morbidity and mortality rates in patients. Hyper induction of pro-inflammatory cytokines, which is known as “cytokine storm”, is closely related to the process of viral infection. However, systemic analyses of H7N9 induced cytokine storm and its relationship with disease progress need further illuminated. In our study we collected 75 samples from 24 clinically confirmed H7N9-infected patients at different time points after hospitalization. Those samples were divided into three groups, which were mild, severe and fatal groups, according to disease severity and final outcome. Human cytokine antibody array was performed to demonstrate the dynamic profile of 80 cytokines and chemokines. By comparison among different prognosis groups and time series, we provide a more comprehensive insight into the hypercytokinemia caused by H7N9 influenza virus infection. Different dynamic changes of cytokines/chemokines were observed in H7N9 infected patients with different severity. Further, 33 cytokines or chemokines were found to be correlated with disease development and 11 of them were identified as potential therapeutic targets. Immuno-modulate the cytokine levels of IL-8, IL-10, BLC, MIP-3a, MCP-1, HGF, OPG, OPN, ENA-78, MDC and TGF-β 3 are supposed to be beneficial in curing H7N9 infected patients. Apart from the identification of 35 independent predictors for H7N9 prognosis, we further established a real-time prediction model with multi-cytokine factors for the first time based on maximal relevance minimal redundancy method, and this model was proved to be powerful in predicting whether the H7N9 infection was severe or fatal. It exhibited promising application in prognosing the outcome of a H7N9 infected patients and thus help doctors take effective treatment strategies accordingly.

## Introduction

Since the first case of human H7N9 influenza virus infection reported in March, 2013 in China, the number increased to about 1564 at the end of Oct, 2017 according to WHO [1]. With a high mortality rate at approximately of 40%, H7N9 virus poses great threat to public health of human beings [2]. Most of the H7N9 influenza virus infected people suffered from severe pneumonia and acute respiratory distress syndrome, and around 63% patients were admitted to an intensive care unit (ICU), about 62% patients had undergone mechanical ventilation [3].

H7N9 influenza virus was better adapted to infect and replicate in human upper and lower airway tissues compared with many other AIVs which was revealed by in vitro studies [4, 5]. It was reported that H7N9 virus could infect higher percentage of human peripheral blood mononuclear cells (PBMCs) than avian influenza A H5N1 virus and pandemic H1N1 virus do [6]. The avian influenza H7N9 virus can even infect BALB/C mice without prior adaption and lead to sever pneumonia which was similar with that in clinical cases [7]. Respiratory epithelia and other types of cells such as alveolar macrophages could be infected by H7N9 virus and die from apoptosis or necrosis followed by induction and release of pro-inflammatory cytokines [8]. Hyper-induction of pro-inflammatory cytokines were well documented in severe H7N9 infected cases and the so called “cytokine storm” showed significant influence on the final outcome of infections [9].

To illuminate the relationship between hyper-induction of cytokines/chemokines and the progression of H7N9 infection, persistent efforts were made. One study found the serum levels of interleukin 8 (IL-8), interferon gamma-induced protein 10 (IP-10), interferon (IFN)-α, IFN-γ, macrophage inflammatory protein 1 alpha (MIP-1α), MIP-1β, monocyte chemotactic protein 1 (MCP-1), and monokine induced by gamma interferon (MIG) were significantly higher in H7N9 virus infected people compared with healthy controls [10]. Another study suggested blood monitoring of IL-8 and IL-6 may helpful in making effective management of severe H7N9 infection cases as the two cytokines were found extremely elevated in patients who died from the infection compared with the discharged counterparts [11]. In a study about 5 hospitalized patients confirmed with H7N9 virus infection, researchers detected high concentration of IP-10, MCP-1, MIG, MIP-1α/β, IL-1β and IL-8 in both sera and broncho-alveolar fluid (BALF) samples. What’s more, they found a positive correlation between high levels of pro-inflammatory cytokines and the severity of clinical outcomes [12].

Here in our study, we systemically investigated the profile of 80 pro-inflammatory cytokines/chemokines in H7N9 virus infected patients at different time points. By grouping patients according to disease severity, we acquired 8 cytokines/chemokines closely related to outcome prediction. Through algorithm based on maximal relevance minimal redundancy method, we established a real-time prediction model to prognose the outcome of H7N9 infected patients. Hopefully the model could be used to assistant in clinical diagnosis and as guidelines for adopting immunomodulatory therapies.

## Results

### Distinguishable cytokine change pattern in clinically confirmed H7N9 infected patients with different severity

Compared with healthy controls, there were 32 cytokines elevated in mild infection group, 39 cytokines elevated in severe infection group, and 35 cytokines elevated in fatal infection group with statistical significance (**Figure 1, Figure 2a**). In total, there were up to 47 of 80 tested cytokines highly expressed in H7N9 infected patients, either mild, severe or fatal patients. Among these 47 cytokines, 24 highly elevated cytokines (IL-13, IL-1a, IL-8, IL-10, IL-7, IL-6; IFN-γ; CSF2, CSF3; BLC, MIP-1b, MCP-2, IP-10, NAP-2, GRO; EGF, TGF-β1, PDGF-BB, IGFBP-2, SCF, HGF, ANG; TNF-a) were shared by all patient groups (**Figure 2a**). While 7 cytokines (IL-12, p70, PARC, TIMP-2, GRO-a, IL-2, TARC, TNF-β) were specifically elevated in severe infection group, 4 cytokines (MIP-1d, MIF, Leptin, IGFBP-1) were exclusively elevated in fatal cases and 1 cytokine (ENA-78) was only detected to be induced in mild infection patients (**Figure 2a**). The results indicated hyper-cytokinemia was induced in patients infected by H7N9 virus and cytokine profile could be different in H7N9 infected people according to severity of the disease.

**Figure 1.**
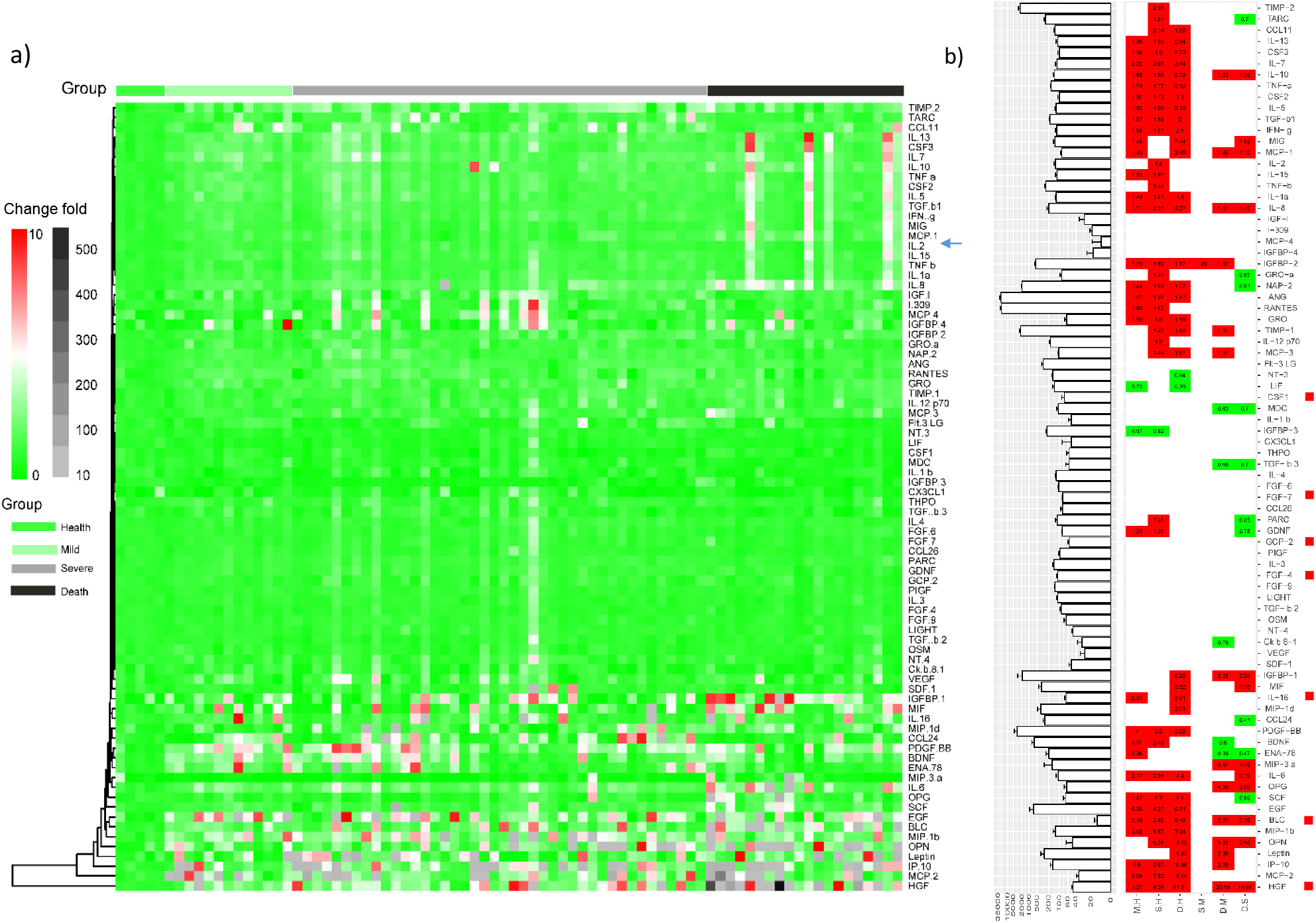
Cytokine expressing change profile during H7N9 infection disease progression. (a) Heatmap of cytokine change fold compared with health individuals. The data was presented as fold of change compared with the mean value of five health controls. Each column represented one patient while each row showed one cytokine or chemocline detected in the study. (b) Different cytokine expression profiles among heathy (H), mild (M), severe (S) and fatal (D) groups. Cytokines sequentially from top to bottom were in accordance with that in Figure 1a. Cytokine expression baselines were indicated by bar chart on the left and the relative fold of change in cytokines or chemokines between different groups was calculated on the right panel. Paired t-test were performed among H, M, S and D. Cytokines or chemokines that up-regulated significantly were indicated in red while those dramatically down-regulated cytokines or chemokines were indicated in green. The seven cytokines or chemokines that showed positive correlation with viral load were highlighted by red solid square.

**Figure 2.**
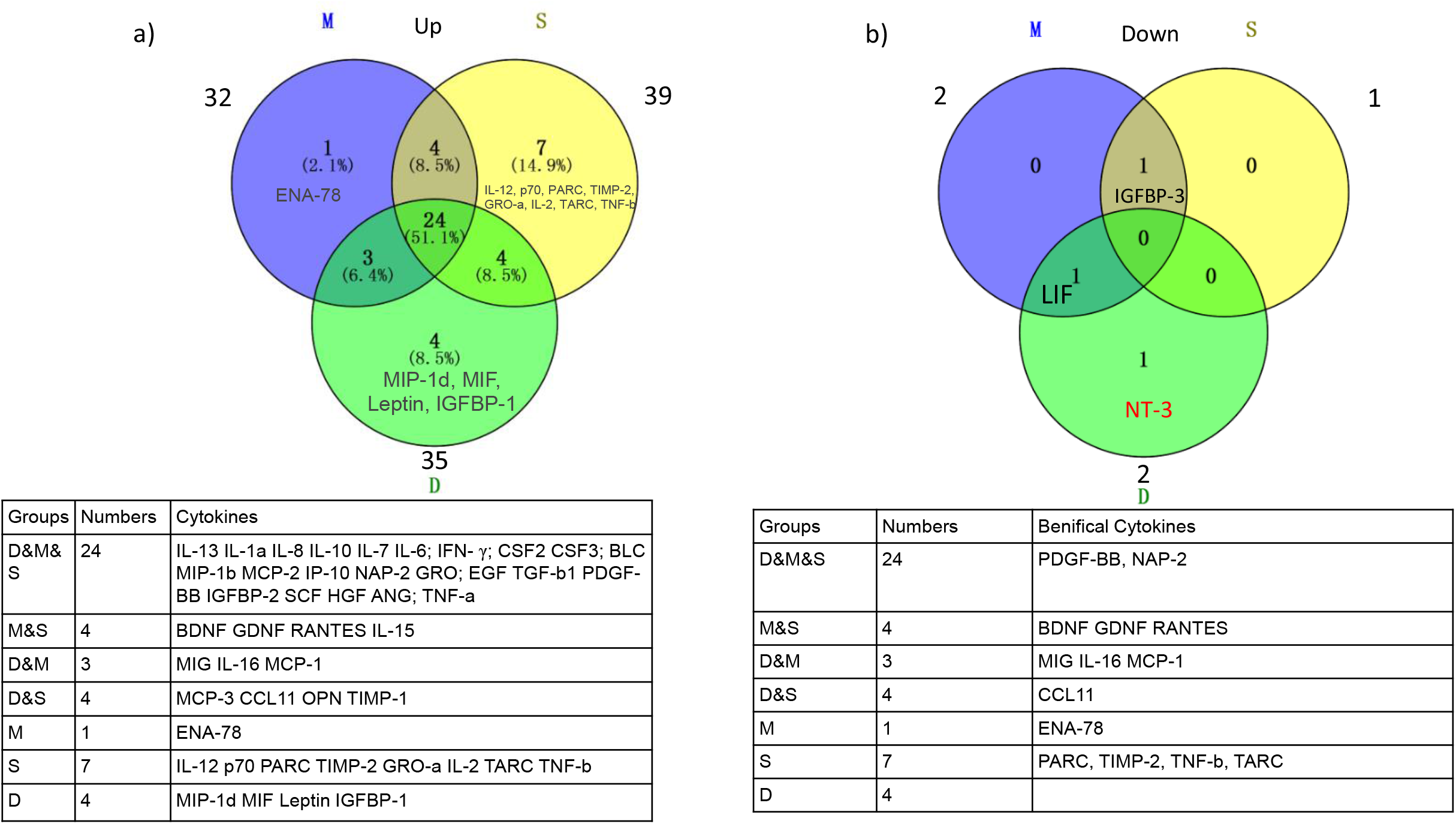
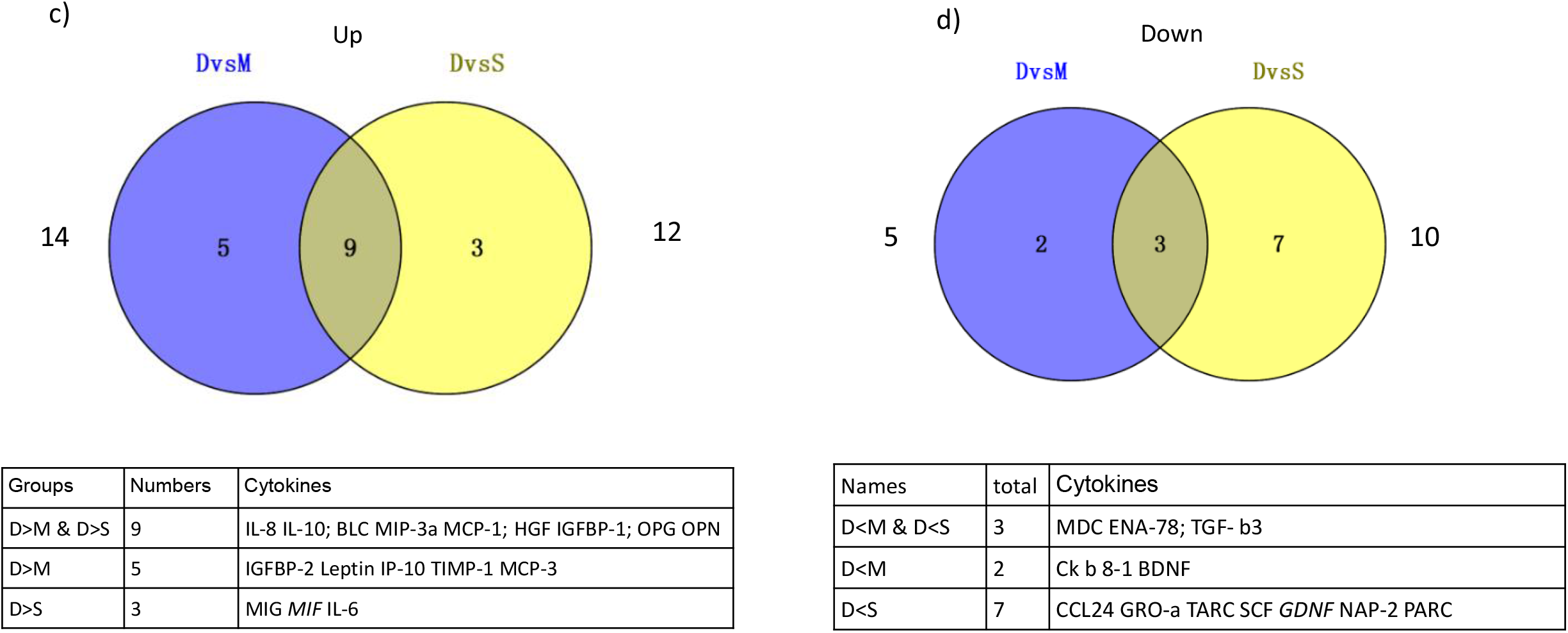
Profiles of cytokine and chemokine response against H7N9 infection among M, S and D patient groups. (a) Venn diagrams of up-regulated and (b) down-regulated cytokines or chemokines in M, S and D groups compared with that in healthy controls. (c) Up-regulated and (d) down-regulated cytokines or chemokines in D group compared with that in M and S group respectively. The exact names of cytokines or chemokines were listed below each venn diagram accordingly.

In contrast to the generally belief that cytokines would be greatly up-regulated during the hyper-cytokinemia caused by H7N9 infections [13], we found several cytokines down-regulated in H7N9-infected patients compared with controls. To be specific, IGFBP-3 and LIF were down-regulated in mild group, IGFBP-3 was down-regulated in severe group, while LIF and NT-3 were down-regulated in fatal group (**Figure 2b**).

### Fatal group showing more intense hypercytokinemia than the mild and severe patient groups

There were 12 and 15 cytokines up-regulated for more than 2 folds in mild and severe groups compared with healthy controls respectively, while amount to 26 cytokines showed more than 2-fold induction in the fatal group. It was notable that five cytokines were changed even more than 10 folds in the fatal group which were 67.21 folds for HGF, 18.44 folds for MCP-2, 16.26 folds for IP-10, 13.42 folds for OPN, and 10.22 folds for Leptin (**Figure 1b**, and **Supplementary Figure 4**).

Compared with mild group, only IGFBP-1 was highly elevated in severe group which indicated that mild and severe outcome patients showed a similar hypercytokinemia pattern. In contrast, fatal patient group showed more cytokines were dysregulated compared with non-fatal groups at different time points. Compared with mild and severe groups, 9 cytokines, which were IL-8, IL-10, BLC, MIP-3a, MCP-1, HGF, IGFBP-1, OPG, and OPN, were significantly elevated in fatal group (P<0.05, **Figure 2c**). What’s more, the 9 cytokines kept at high levels through-out the whole course of viral infection (**Figure 3a**). Additionally, there were three cytokines, MDC, ENA-78 and TGF-β3 were significantly down-regulated in the fatal group compared with the other two infection groups (**Figure 2d**), and ENA-78 as well as MDC and TGF-β3 showed a persistent trend at lower levels at different time points (**Figure 3b**). The three down-regulated cytokines may play key roles through-out the course of H7N9 infections (See below analysis).

**Figure 3.**
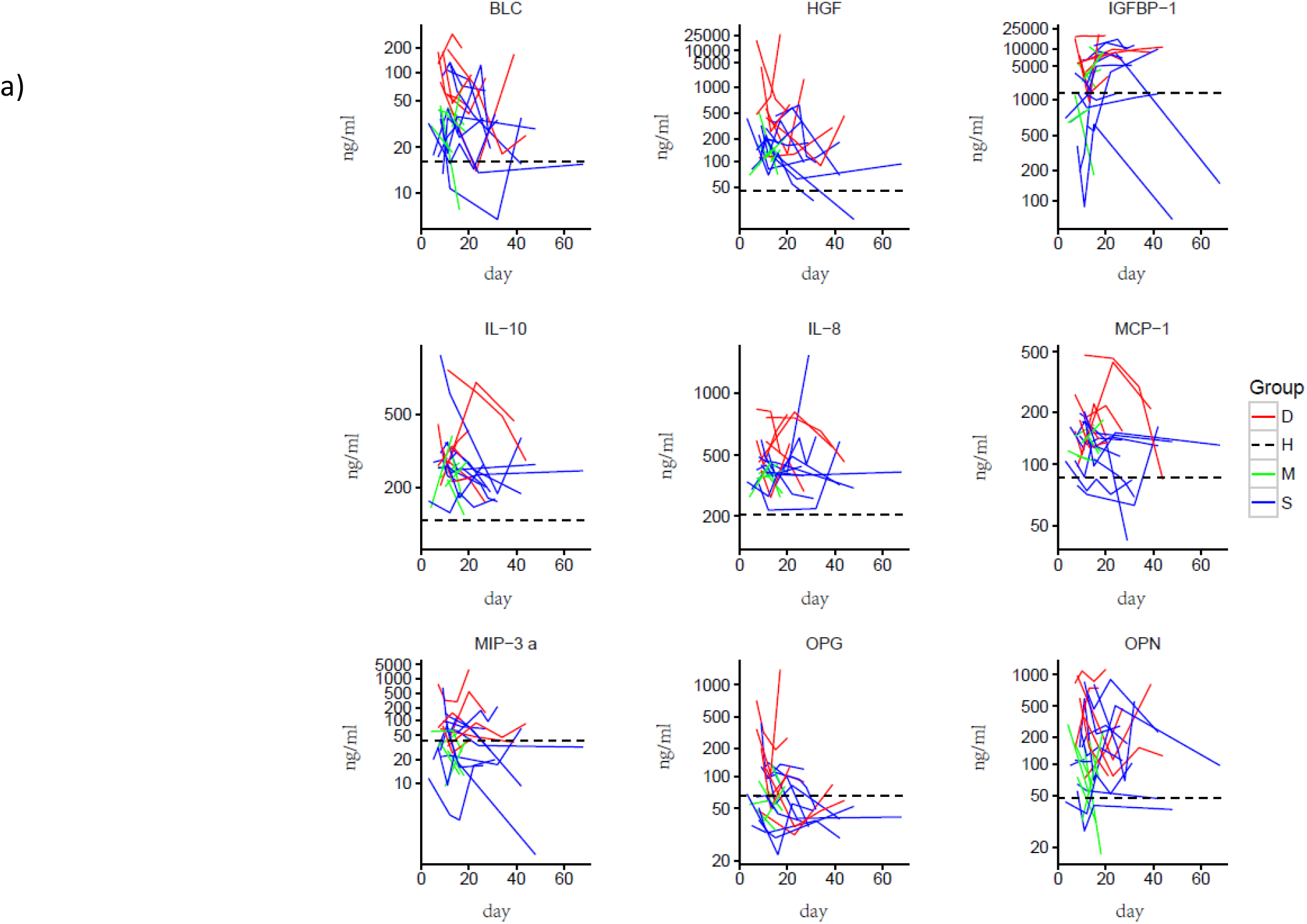
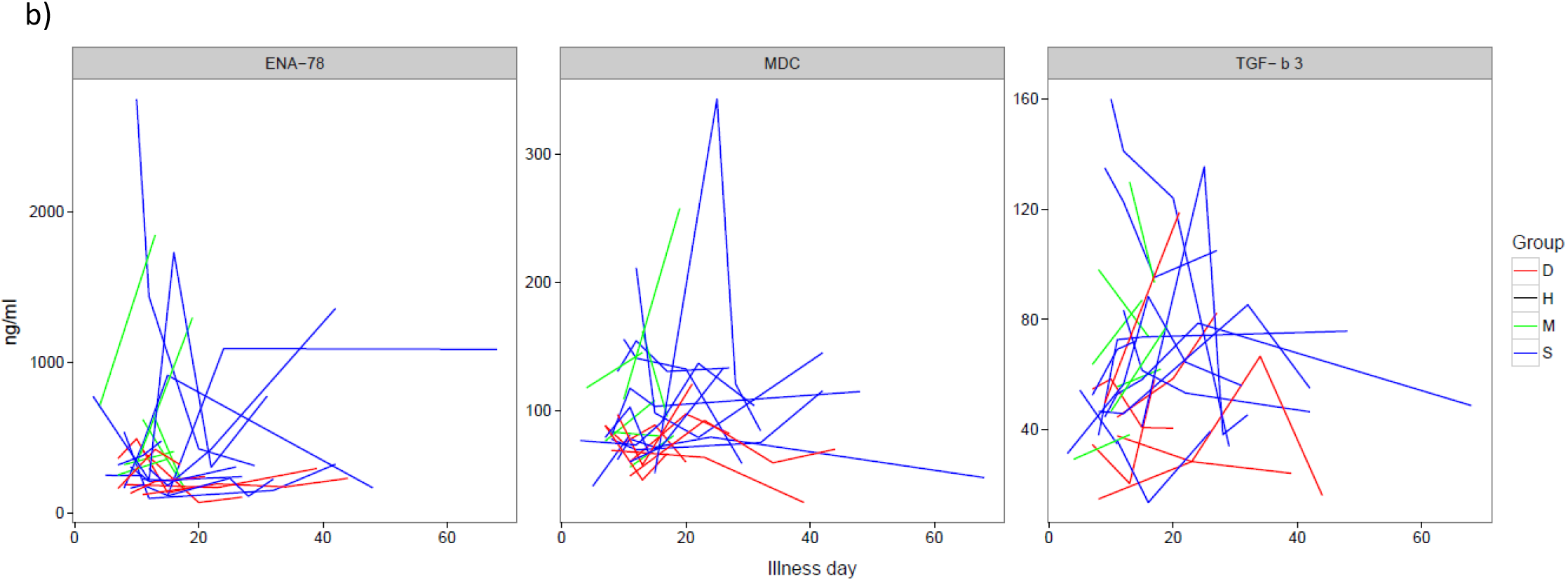
Different Kinetics of dys-regulated cytokines or chemokines in fatal-outcome patients. (a) Dynamic kinetics of cytokines or chemokines upregulated or (b) downregulated significantly in fatal patients compared with both mild and severe patients.

### Various cytokines correlated to clinical manifestation of H7N9 infection disease but dysregulated in fatal group patients

To reveal the relation between cytokine regulation and manifestations of patients after H7N9 viral infection, and to explain the contributions of the dysregulated cytokines to hypercytokinemia syndrome, a serial of correlation analyses was performed. Based on spearman’s rank correlation analysis, 42 cytokines were found correlated with at least one of the following clinical manifestations which were C-reactive protein (CRP), oxygenation index (OI), procalcitonin (PCT), and body temperature (T). Among the 42 cytokines, 21 of them showed negative correlation with CRP and/or PCT and/or T, and/or showed positive correlation with OI values (**Figure 4**). Elevation of the 21 cytokines were supposed to be beneficial for the prognosis. Similarly, we identified 14 cytokines whose elevation were related to more severe symptoms, which are positively correlated with CRP or/and PCT or/and body temperature, or/and negatively correlated with OI values (**Figure 4**).

**Figure 4.**
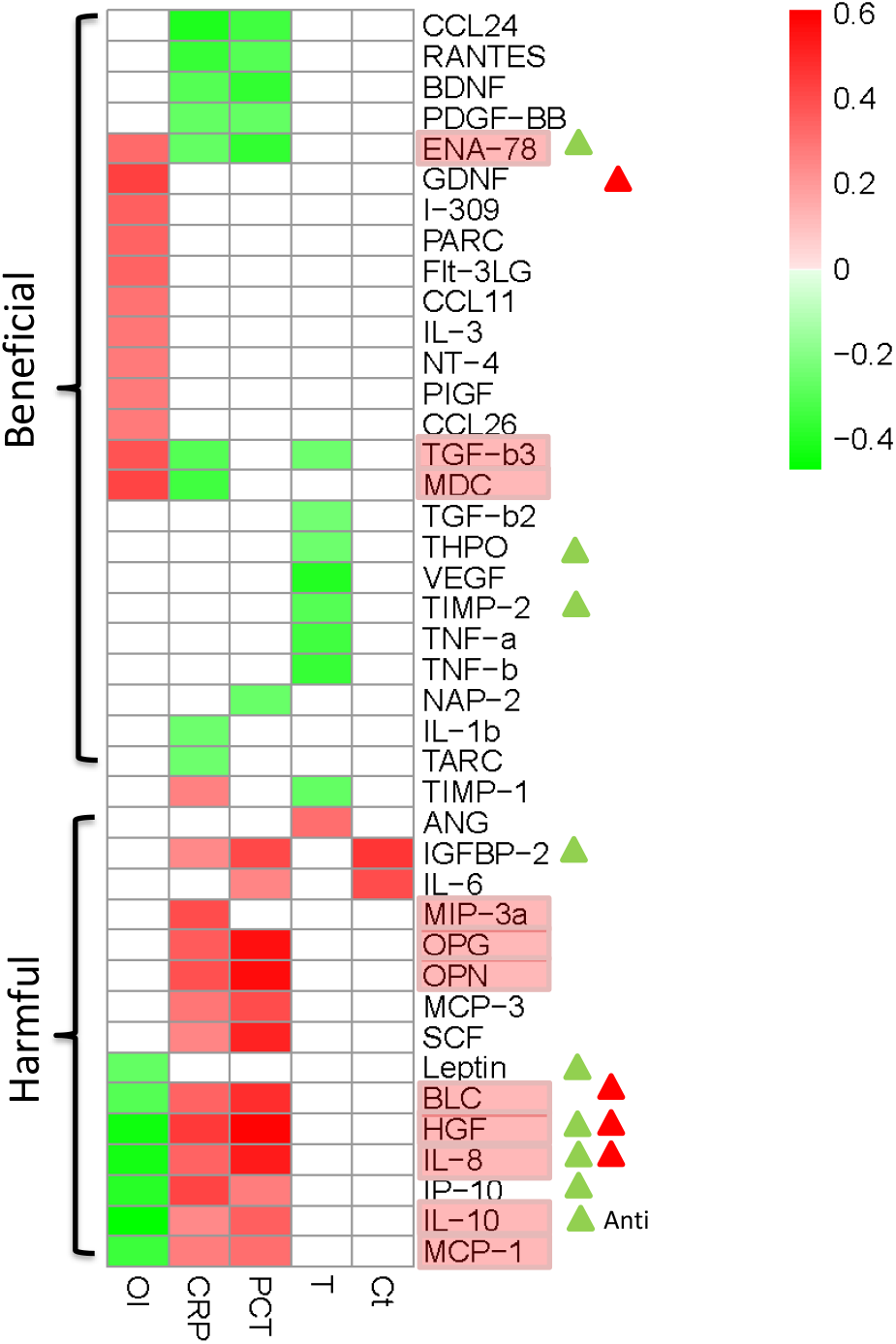
Correlation analysis of cytokine or chemokine levels with clinical manifestations of H7N9-infected patients. Spearman’s rank correlation analysis was performed and 42 cytokines or chemokines showed correlations with at least one of the five clinical manifestations which were OI, CRP, PCT, T (body temperature) and viral load (shown as CT value). The correlation was evaluated based on correlation coefficient (r) value as r=0.2-0.4 for weak correlation, r=0.4-0.6 for moderate correlation and r=0.6-0.8 for strong correlation.

Surprisingly, in fatal patients, we found all 9 specific highly elevated cytokines except IGFBP-1, are harmful, and 3 cytokines, specific down-regulated in fatal patient, were beneficial when elevated. In the fatal group patients, MDC (0.63 and 0.70 fold to that of mild and severe patient groups respectively), ENA-78 (0.39 and 0.47) and TGF-β3 (0.68 and 0.70) were expressed at lower level (**Figure 1b** and **Supplemental Figure 3b**). Especially the MDC and ENA-78 were kept at lower level during whole hospitalization (**Figure 3b**). These results suggested that the expression of MDC, ENA-78, TGF-β3 had been suppressed or those related pathways may not being activated efficiently. Besides, 6 elevating-beneficial cytokines, CCL24, TARC, GDNF, PARC, BDNF and NAP-2 were deficiently responsed in fatal group compared with mild or severe group while the 4 elevating-harmful cytokines, IGFBP-2, IL-6, IP-10, Leptin and TIMP-1 were elevated in fatal group patients compare with mild or severe groups ones (**Figure 4** and **Supplementary Figure 4**). These results indicated a distinct pro-inflammatory cytokine induction profile in fatal H7N9 infected patients comparing with mild and severe counterparts, and that mounting 21 cytokines, were abnormal and doing harm, as mediators, to fatal group patients (**Figure 1b** and **4**; **Supplemental Figure 4**). In contrast, these cytokines were at moderated levels in all mild and recovery patients, which is supposed to be under control of host immune system. Furthermore, it is reasonable to speculate that drugs or therapies helping for elevating these three cytokines, especially MDC, ENA-78, TGF-β3, and targeting for decreasing the 8 major harmful cytokines, would be benefit for survival and recovery in H7N9 infection.

### Cytokine co-regulation network during disease progress: dysregulation of cytokines in fatal patients

To analyze the correlation between those cytokines and chemokines during hyper-cytokinemia, spearman’s rank correlation analysis was performed pair by pair, and cytokine network charts were mapped separately according to the severity of H7N9 viral infection. In those charts, a total of 1372 pairs of cytokines or chemokines were significantly correlated (r>0.8 or r<−0.8, P<0.05) and line-linked (**Figure 5**). Compared with healthy and mild samples (**Figure 5a, b**), severe and fatal samples at early time-points formed networks with cluster centers (**Figure 5c, e**). However, the cluster centers remain in fatal cases even at late infection stage (**Figure 5f**) while the clusters disappeared and the correlations between cytokines became weak in severe cases before discharged from hospital (**Figure 5d**). It was worth noting that cytokine network of severe cases at early time-point was characterized by single cluster core which may indicate a more coordinating regulation of inflammation after H7N9 infection in severe patients compared with that in fatal patients as there showed generally two cluster cores in their cytokine networks. The lack of cytokine coordination in fatal cases was also implied by the increased frequencies of negative correlations between cytokines within cluster cores.

**Figure 5.**
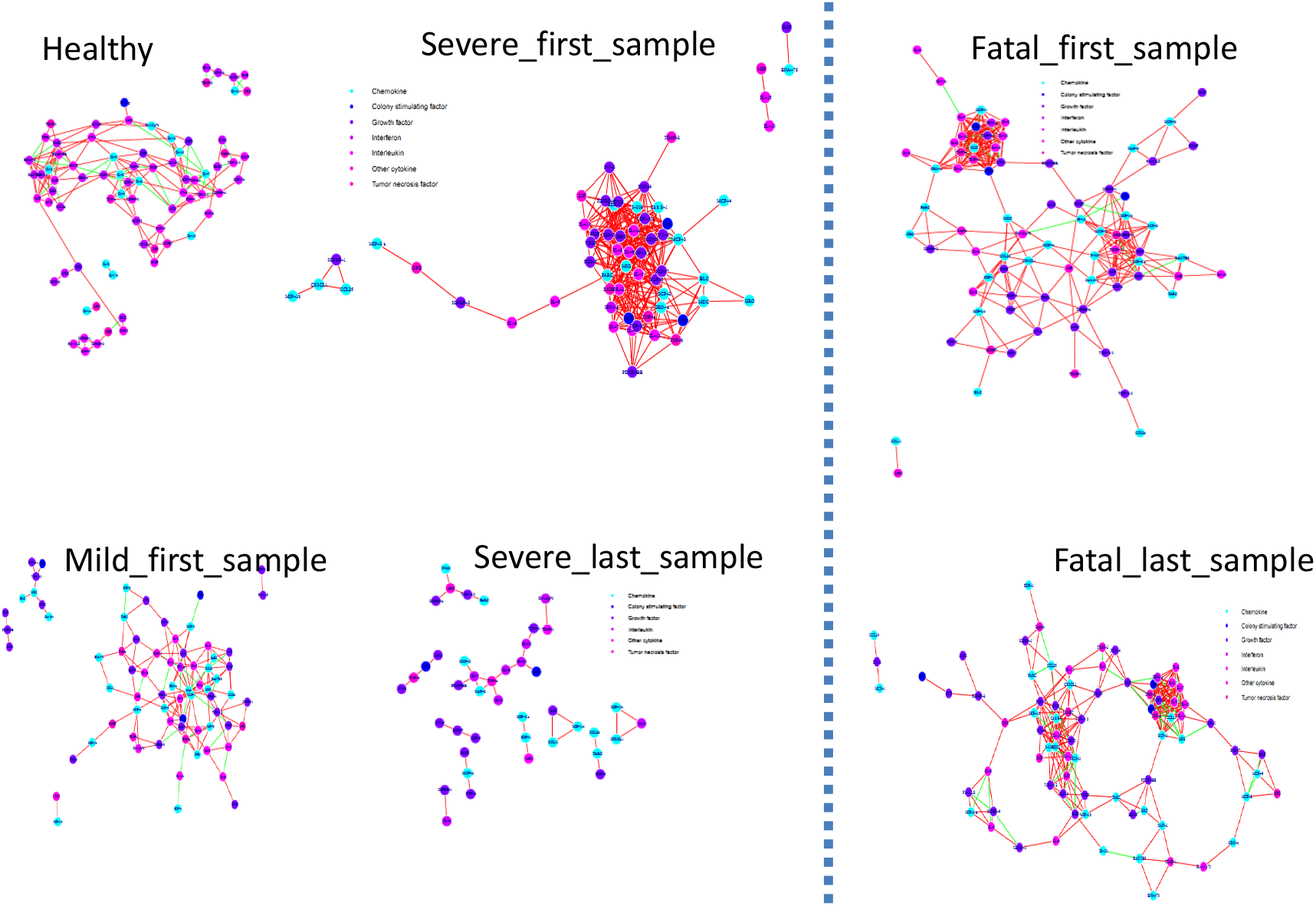
Co-regulation networks of cytokines and chemokines in H7N9 infected patients and healthy controls. Cytokines or chemokines were shown in different colours according to their properties. The links indicated strong correlations between each two cytokines/chemokines (r>0.8) according to spearman’s rank correlation analysis.

### Real time predicting model establishing based on cytokines profiles

As reported, cytokine levels could be used as predicative indicators of clinical outcomes in patients identified with H7N9 virus infection [14]. To evaluate the potential prediction ability of each cytokine, we analyzed the 80 cytokines or chemokines one by one based on receiver operating characteristic (ROC) curve method. The area values under the ROC curve (AUC) of each cytokine was represented in a heat map for predicting illness (Fatal+Severe+Mild groups vs Healthy), Fatality (Fatal group vs Severe+Mild groups), Mild (Fatal+Severe groups vs Mild group) as well as recovery (Severe-first vs Severe-last time) (**Figure 6**). 35 cytokines showed values of AUC ≥0.8, which suggested they could be used for prognosis of H7N9 virus infection with an accuracy of no less than 80%. BLC was found to be able to predict 94% fatality cases, higher than all previous reported cytokine biomarkers, which highlight the importance of this cytokine during H7N9 virus infection for the first time. In addition, MIF (82%), IGFBP1 (82%), HGF (86%) and IL-8 (89%) were all promising predictors for fatality cases. For mild cases, IL-10 (83%) as well as IL-8 (82%) turned out to be good predictors while only Flt-3-LG (84%) showed the potential to predict recovery from severe cases (**Supplementary Figure 7**).

**Figure 6.**
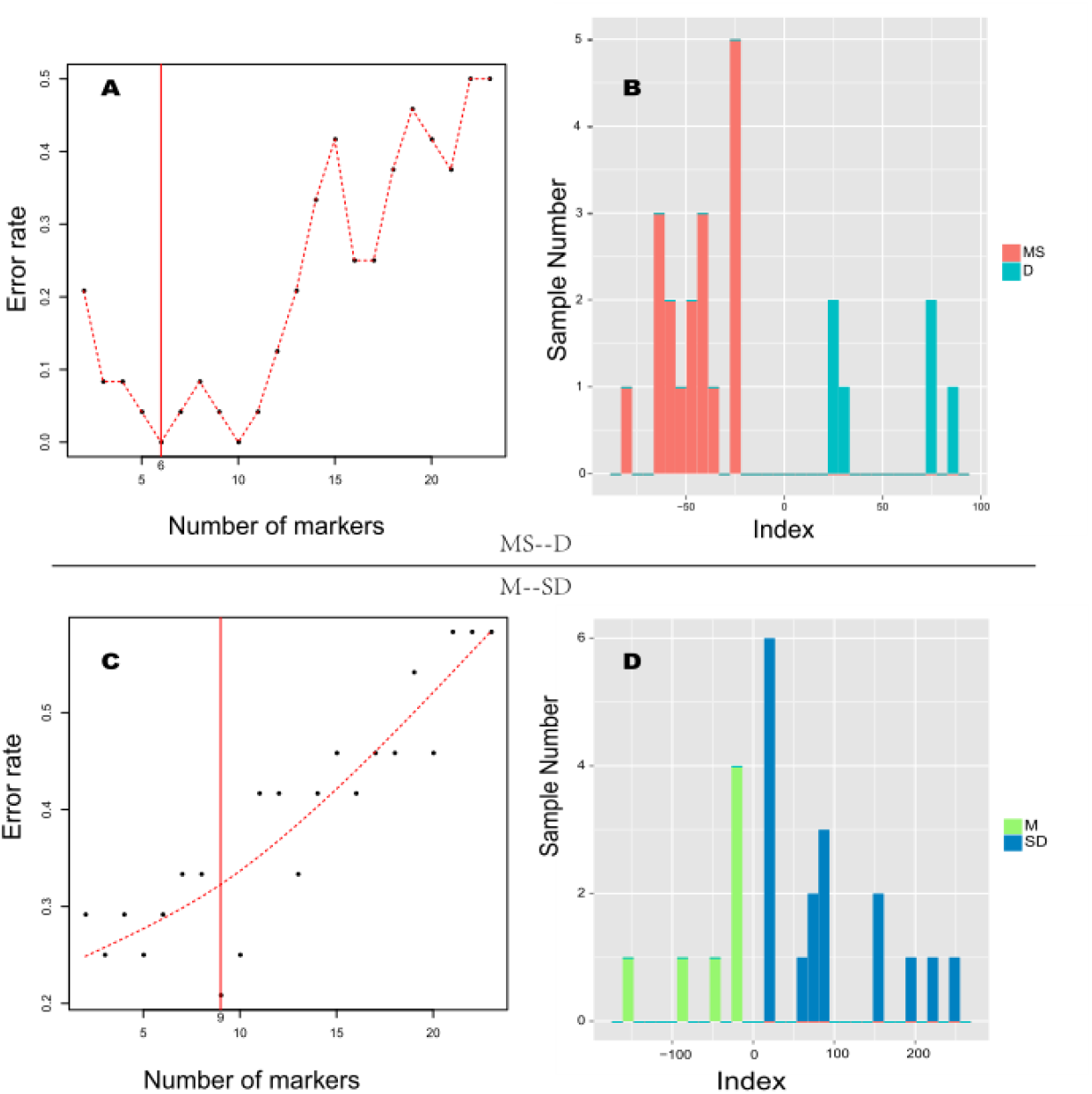
Establishment of prediction models. The formulas were inducted by minimum redundancy maximum relevance (mRMR) algorithm and the induction incorporated only the firstly sampling data of cytokine/chemokine levels in H7N9 infected patients. (a) Lowest error rate for factor selection for distinguishing Mild and Severe (MS) vs fatal (D). (b) Index values for distinguish Mild and Severe (MS) vs fatal (D) of each sample. (c) Lowest error rate for factor selection for distinguishing Mild (M) vs Severe and fatal (SD). (d) Index values for distinguish Mild (M) vs Severe and fatal (SD).

It is assumed that power of prognosis predicting would be increased using the different clinical features of multi-cytokines. It is also plausible to test for real time prognosis assessment using the multi-cytokines model. To construct a multi-cytokine prognosis predication model, the minimum redundancy maximum relevance (mRMR) algorithm was applied to the first time-point samples of all patients in our dataset, which containing 7 samples of mild patients, 11 samples of severe patients and 6 samples of fatal patients.

In case of predicting mild outcomes in people after infection of H7N9 virus, we divided the data into two groups which was the mild patients group (M) and the severe plus the fatal patients (S+D) group. Based on mRMR sorting and selection results, it was found a minimal usage of concentrations of 9 cytokines could distinguish M group from S+D group with an error rate at 0 (**Figure 6**). The mathematical model was displayed as below and the prognosis would be mild if the index value was negative:

### M-SD index value = −211.6204 + (0.2094 * IL8 + 0.1978 * IGFBP2+ 0.0376 * Leptin + 0.0323 * ENA78 + 0.0136 * IP10 + 0.0020 * TIMP2) – (1.381 * THPO + 0.0808 * IL10 + 0.0046 * HGF)

Similarly, to predict the fatal outcomes, we divided the data into mild and severe (M+S) group and fatal (D) group. Six cytokines were selected and their concentrations could be used to distinguish D group from M+S group with an error rate at 0.2 (**Figure 6**). The formula was displayed as below and positive index values indicated a fatal outcome.

### MS-D Index value = 6.9210 + (0.2686 * BLC + 0.0809 *IL8 + 0.004*IGFBP1 + 0.0107 * MIF) – (1.0847 * GDNF + 0.0017* HGF)

The two indices efficiently divided the samples of first time point of three symptom groups in a 2-D coordinate, Mild in Quadrant-III (M-SD index<0 & MS-D index<0), Severe in Quadrant-IV (M-SD index>0 & MS-D index<0) and Fatal in Quadrant-I (M-SD index >0). (Figure 8A). To validate the two indices, later time-point samples and healthy samples were applied to the two formulas. All healthy and most of the mild samples (10 of 11 samples) were retained in Quadrant-III at different time points (**Figure 7B**). Interestingly, samples of the severe group cases displayed a noticeable trend of moving from Quadrant-IV towards the Healthy and Mild section (Quadrant-III), and few time points of single patient entering the Fatal Quadrant I (only two time points of two patients, S1 and S8), demonstrating recovery during treatment (**Figure 7C**). The Fatal samples mainly remained in Quadrant-I, again demonstrating the robustness of the indexes (**Figure 7D**). For summarize, we established a two-dimensional coordinate system and a combining of 13 cytokines could assessment disease status and predict prognosis precisely.

**Figure 7.**
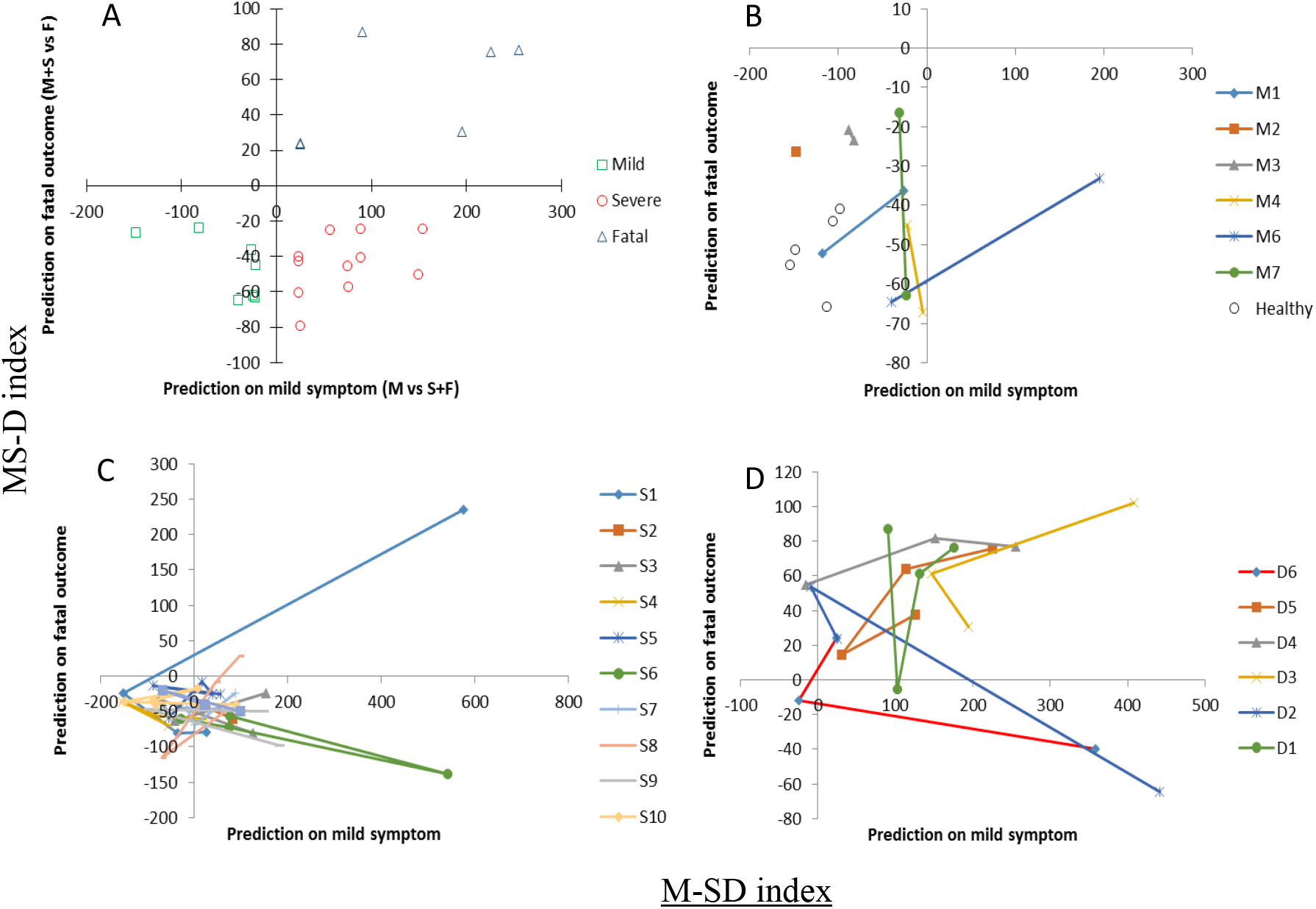
Validation and proof of prediction models. (a) Training dataset (1^st^ sample) (b) Validation with all samples of healthy and mild symptom. (c) Validation with all samples of fatal outcome. (d) Validation with all samples of severe symptom.

## Discussion

80 cytokines and chemokines profiles were investigated in patients infected with H7N9 virus. This was the most comprehensive study as far as we know and gave a panorama of hypercytokinemia in patients with different severity. It has been reported that IL-2, IL-6, IL-10, IL-17, IL-18, IFN-γ, TNF-α, MIF, SCF, MCP-1, HGF, SCGF-β and IP-10 were significantly elevated in H7N9 virus infected patients [14–17]. However, here we revealed up to 24 cytokines and chemokines were up-regulated remarkably in infected cases compared with healthy controls during H7N9 viral infection and many of them were reported for the first time yet showed great induction on protein levels.

For example, compared with healthy people, BLC elevated for 2.19, 2.69 and 6.43 folds in mild, severe and fatal cases on protein levels respectively. Besides, the protein levels of MCP-2 increased 8.56, 7.92 and 9.44 folds in mild, severe and fatal cases comparing with basal levels. Both of the inflammatory factors were chemokines with BLC targeted at B cells while MCP-2 could activate many kinds of immune cells such as master cells, monocytes, NK cells and T cells [18–20]. Our findings, especially the discovery of novel cytokines and chemokines that presented remarkable changes in people infected with H7N9 virus, provided new targets for adopting immuno-modulatory therapies and gave clues for illuminating mechanisms of hypercytokinemia.

Great attention was paid to the highly induced cytokines or chemokines after H7N9 viral infection while few people talked about those downregulated cytokines or chemokines [21, 22]. In an *in vitro* cell culture model infected with H7N9 virus, nine cytokines were reported to be down-regulated by 2 folds compared with clean controls [23]. However, the cytokine responses in cell cultures may not reflex the real scenario in patients infected with H7N9 virus. Our study of clinical cases showed there were 3 cytokines and chemokines down-regulated significantly and none of them in accordance with the previous report. For the three cytokines and chemokines, IGFBP-3 was firstly recognized as bioavailability regulator of insulin-like growth factors (IGFs) which could regulate the entry of IGFs to tissues [24]. However, later studies showed IGFBP-3 played more roles. For instance, through binding to different ligands, IGFBP-3 could induce apoptosis alone or in conjunction with certain agents, or contribute to the repair of damaged DNA [25]. LIF could prompt the differentiation of myeloid leukemia cells and showed anti-inflammation properties [26, 27]. NT-3, which is a neurotrophic factor in the nerve growth factor family, was also detected in monocytes and play roles in immune responses [28]. Based on the properties of the three cytokines or chemokines, further study may shed light on the underlying mechanism of H7N9-induced cell death, dysregulation of inflammatory response and impaired antibody production after H7N9 viral infections [29, 30].

It was proved that patients with severe influenza infections were more likely to have higher temperature and viral load [31], lower CRP levels [32], decreased OI [33], as well as lower PCT levels [34]. In this study we correlated the cytokine and chemokine levels with these parameters innovatively and distinguished those pro-inflammatory factors according to their beneficial or detrimental effects on clinical outcomes. ENA-78, IP-10, IL-8, TIMP-2, IGFBP-2, Leptin, IL-10, THPO, HGF, BLC, MIF, IGFBP-1, and GDNF were especially emphasized here as the 13 cytokines and chemokines showed predictive potential in H7N9 infected cases. Among them, IP-10, HGF, MIF were found to be independent outcome predictors in H7N9 virus infected patients previously which was in accordance with our findings [35]. However, we further explored the possibility of establishing a method that could make real-time prediction of clinical outcomes after H7N9 infection based on as less as those cytokines or chemokines. Though there were data showed IL-6 >97 pg/mL, IL-8 >40 pg/mL and CRP >90 mg/L in serum may indicate adverse clinical outcomes, or studies declared a significant association between CRP level and fatality outcome, they did not possess the ability of making real-time prediction in H7N9 infected patients [35–37]. The multi-factor prediction models we established in the study turned out to be powerful and accurate as revealed by the verification data. It would be hopefully that the formulas could be applied to help in precise treatment in H7N9 infected patients, through getting a preliminary judgement on the prognosis of a H7N9 virus infected patient as early as possible, and clearly that is vital for later following up management.

## Methods

### Patients and sample collection

The study protocol was approved by the Institutional Review Board of BGI IRB consent and Peking university Shenzhen hospital IRB. Informed written consent was obtained from all participants. 18 nonfatal H7N9-infected patients (7 mild patients, 11 severe patients), 6 fatal H7N9-infected patients and 5 healthy controls were enrolled in the study (**Figure S1a**). All patients with H7N9 infection were confirmed by real-time PCR and were admitted to the Shenzhen Third People’s Hospital. There showed no statistical difference in terms of age between fatal patients and no-fatal patient (mean±SD. 50.3±20.1 vs 54.2±16.0). The duration of hospitalization was longer in fatal group than nonfatal group (mean±SD. 18.6±9.9 vs 36.3±6.9, *P*<0.0001). Most of the patients were admitted to hospital within 10 days after symptom onset and were given anti-viral treatment immediate after admission. Other clinical characteristics were similar between two groups (data not shown).

Patients were divided into mild group, severe group and death group according to the guidelines for avian influenza A(H7N9) virus [38]. Blood samples were collected on the first day of admission and throughout of the process of the disease. There were 5 samples collected within 7 days after admission and 8 samples collected within 7 days before discharge in mild patient group (mild group). For severe infection (severe group) and deadly cases (fatal group), 8 and 5 samples were collected within 7 days after admission, 25 and 13 samples were collected during illness progression stage, 9 and 2 samples were collected either within 7 days before discharge or death, respectively (**Supplemental Figure 1a**).

### Viral RNA extraction and real-time PCR

RNA was extracted from the samples by QIAamp viral RNA mini kit. Standard Real-time reverse transcription polymerase chain reaction (RT-PCR) assay for H7N9 confirmation was performed in Shenzhen Center for Disease Control (CDC). The results of positive or negative were judged according to the Guidelines authorized by China National Influenza Center of China CDC [38].

### Serum Cytokine and chemokine arrays

EDTA-anticoagulant tubes were used to collect blood samples. Plasma was separated by centrifugation (3000g for 10 min) at 4°C and stored in −80°C until analysis. We analyzed 80 cytokines and chemokines (14 interleukins; 1 interferon; 4 tumor necrosis factors; 3 colony stimulating factors; 26 growth factors; 26 chemokines; 6 other cytokines; Table S2) with the RayBio^®^ Human Cytokine Antibody Array 5(G-Series) according to the manufacturers’ instructions.

### Statistical analysis and co-regulation network constructions

Wilcoxon signed-rank test was used to determine whether the differences between two groups were statistical significance. Spearman’s rank correlation coefficient was adopted to analyze the line correlation. Any value of *P* <0.05 was considered statistically significant. Statistics and plotting were done using libraries implanted in Rstudio version 1.1.

### ROC predicting analysis

We calculated the receiver operating characteristics (ROC) curves for outcome predicting analysis using single cytokine.

### mRMR predict model establishment

All of the samples were classed into 2 compare group: M vs SD (mild as one group, severe and death as one group), MS vs D (mild and severe as one group, death as one group). The minimum redundancy maximum relevance (mRMR) feature selection method described by Peng *et al* [39] was used to calculate the redundancy coefficient for each cytokine between group respectively in M vs SD and MS vs D, which was used to sort the cytokine.

The accuracy of each model was evaluated by leave-one-out cross-validation (LOOCV) to find the optimum subset for building a linear discrimination classifier. We chose the lowest error rate model as the final model to predict the remaining samples such as healthy control, mild and severe samples in different time points. Two of the linear regression formula, respectively for 2 compare group, was as follows:

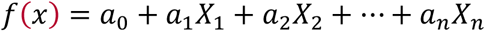

where “*X_i_*” refers the expression of the cytokine selected from mRMR selection and *a_i_* indicates the redundancy coefficient of each cytokine.

## Ethical approval

All procedures performed in the studies involving human participants were approved in accordance with the ethical standards of the institutional and/or national research committee and with the 1964 Helsinki declaration and its later amendments or comparable ethical standards.

## Acknowledgements

This study was supported by the Shenzhen Science and Technology Research and Development projects (JCYJ20150402111430617 and JCYJ20160427151920801).

